# An immunocompetent model of MCPyV-driven Merkel cell carcinoma reveals tumor evolution under immune selection

**DOI:** 10.64898/2026.07.10.737822

**Authors:** James M Regan, Xiaomei Li, Mykaela Salvacion, Tiana Luo, Meng Jia, Grace Ho, Jonathan R Xu, Shujing Liu, Zhi Huang, Xiaowei Xu, Jianxin You

## Abstract

Merkel cell carcinoma (MCC) is a neuroendocrine skin tumor that is frequently driven by integration of Merkel cell polyomavirus (MCPyV). In MCC, the MCPyV genome is truncated, but expression of the viral tumor antigens, truncated large tumor antigen (LTT) and small tumor antigen (sT), is maintained and drives uncontrolled proliferation. We introduced constitutive expression of the MCPyV T antigens (TAs) into primary mouse dermal fibroblasts (MDFs) to determine whether these cells are susceptible to MCPyV-driven transformation. TA expression alone in MDFs induced key MCC markers, cytokeratin-20 (CK20) and Sry-box transcription factor 2 (SOX2), and promoted anchorage-independent growth indicative of cellular transformation. Subcutaneous implantation of TA-transformed fibroblasts produced high-grade MCC-like tumors that grew persistently in immunodeficient NSG mice but not in immunocompetent C57BL/6 mice. Serial *in vivo* passaging of the tumor cell line enhanced tumor growth, reduced expression of p53-target genes and MHC-1, and was accompanied by a shift in T antigen isoform expression, with decreased LTT and increased sT expression. Our data demonstrate that MCPyV-driven tumors acquire immune-evasive adaptions during tumor progression *in vivo* and suggest that the anti-tumor immune response exerts selective pressure in MCC that favors expression of sT rather than LTT. The model established in this study provides a unique platform for studying evolution of MCPyV-driven tumors under immune pressure and identifying mechanisms of immune evasion in MCC that could be used to develop new therapeutic strategies.

**Significance Statement:** MCPyV tumor antigen expression transforms mouse dermal fibroblasts to generate MCC-like tumors. Serial *in vivo* passaging reveals tumor evolution under immune pressure, providing a model to study immune evasion mechanisms in MCC.

## Introduction

Merkel cell carcinoma (MCC) is a rare neuroendocrine tumor of the skin with poor clinical outcomes and a rising incidence(1–3). Surgery is the first-line treatment option for local MCC but patients often present with more advanced disease(4). MCC frequently metastasizes to lymph nodes and other distant sites where systemic treatments such as immune checkpoint inhibitor therapy are needed(5). While the PD-L1/PD-1 immune checkpoint has been a successful strategy in treating advanced MCC(5, 6), almost half of all patients do not respond to these treatments(7). Improved experimental models are needed to better understand MCPyV-driven oncogenesis and identify new therapeutic strategies.

In 2008, a small DNA virus, Merkel cell polyomavirus (MCPyV), was found to be integrated into the host genome in about 80% of MCC cases(8). The MCPyV genome has two coding regions, which are expressed at different time points in the viral life cycle. The early region encodes the nonstructural proteins large T antigen (LT), small T antigen (sT), 57kT antigen, and alternative large T antigen open reading frame (ALTO), and the late region encodes the capsid proteins VP1 and VP2. Both regions are under the control of the non-coding regulatory region (NCRR). The MCPyV genome is integrated in MCC resulting in a truncated form of LT (LTT). In MCC with integrated MCPyV (MCPyV+ MCC), the tumor mutational burden is relatively low when compared to MCC without integrated MCPyV (MCPyV- MCC)(9, 10). This contrast suggests that integration of MCPyV is an important factor in oncogenesis of MCPyV+ MCC. Expectedly, expression of the viral T antigens (TAs) is maintained as a key feature of the tumor and is required for continued proliferation(11).

Both LTT and sT possess oncogenic functions and have been demonstrated to have transforming capabilities *in vitro*(12, 13). The major oncogenic function of LTT involves inhibition of the tumor suppressor retinoblastoma protein (Rb). Truncation of the MCPyV genome removes most functional regions of LT but retains an LxCxE motif that is conserved in polyomaviruses(14, 15). In MCPyV+ MCC, the LxCxE motif of LTT binds to Rb and prevents it from repressing E2F transcription, leading to uncontrolled progression through the cell cycle(16, 17). sT has multiple oncogenic functions, impacting both translation and transcription in the host cell. sT has been shown to dysregulate cap-dependent translation by reducing turnover of hyperphosphorylated eukaryotic translation initiation factor 4E-binding protein 1(4E-BP1)(12). Hyperphosphorylation of 4E-BP1 was then demonstrated to be necessary for sT-induced transformation(12). Previous work has also shown that sT can interact with transcription factors MYCL and EP400 to induce transcriptional changes and indirectly inhibit the tumor suppressor tumor protein 53 (p53)(18, 19). Expression of sT has also been shown to modulate the interferon response which may contribute to immune evasion in MCC(20).

Several mouse models of MCC have been created by expressing MCPyV TAs under the control of epithelial promoters, resulting in epithelial hyperplasia and increased cellular proliferation(21, 22), and the development of cutaneous papillomas in some mice(23). Other models combining MCPyV TA expression with additional genetic alterations, including Merkel cell lineage reprograming and loss of tumor suppressors such as p53 or Rb, have achieved tumor growth in mice that closely resembles human MCC(24–27). Most existing MCC mouse models assume an epithelial origin of MCC, but human dermal fibroblasts (HDFs) represent a compelling alternative candidate because they are the only cell type currently known to support the complete MCPyV life cycle *in vitro*(28). Recently, MCPyV DNA was detected in the dermal papillae of healthy skin, suggesting that the reservoir for MCPyV in the skin may reside in the dermal compartment of the hair follicle(29). MCPyV+ MCC lacks the UV mutational signature characteristic of epidermal tumors, suggesting that the cell of origin may reside in a UV-protected niche such as the dermis(30). These observations suggest that dermal fibroblasts could be susceptible to MCPyV-mediated oncogenic transformation. To address this, we investigated whether expression of MCPyV TAs alone is sufficient to transform primary mouse dermal fibroblasts (MDFs). We demonstrate that TA-expressing MDFs undergo oncogenic transformation, exhibiting anchorage-independent growth, expression of canonical MCC markers including cytokeratin-20 (CK20) and sry-box transcription factor 2 (SOX2), transcriptional programs characteristic of MCC, and formation of MCC-like tumors *in vivo*.

Although MCPyV+ MCC is considered highly immunogenic owing to its high expression of viral proteins, the tumor also possesses several immunosuppressive mechanisms(31). Our results show that MCPyV TAs were sufficient to transform MDFs, but the transformation did not confer sufficient immune-evasive properties to support progressive growth in immunocompetent C57BL/6 mice. The inherent immunogenicity of the model provided an opportunity to investigate how MCPyV-driven tumors adapt under host immune pressure. We performed serial *in vivo* passaging of the TA-transformed cells, which selected for improved tumor growth, downregulation of p53 target gene expression and MHC-1 expression, and strikingly, decreased LTT transcription with a simultaneous increase in sT transcription. These results demonstrate that MDFs are permissive to MCPyV-driven transformation and can provide an immunocompetent model for studying MCPyV-driven tumorigenesis and tumor evolution under host-immune pressure.

## Materials and Methods

### Isolation of mouse dermal fibroblasts

Ear tissue was obtained from C57BL/6 mice. Tissue was submerged in 70% ethanol for 10 seconds and air dried for 5 minutes at room temperature. The tissue was cut into several pieces and placed into a solution of Dulbecco’s phosphate buffered saline (DPBS) (Life technologies, 14190-136) containing 10mg/mL Dispase II (Sigma, D4693) and 5X antibiotic-antimycotic (Life technologies, 15240-062), and incubated for 16 hours at 4°C. The tissue was then briefly rinsed in DPBS before being minced and incubated in a solution of DMEM/F12 (Life technologies, 113200-33) containing 1.5mg/mL Collagenase IV (Life technologies, 17104-019) and 5X antibiotic-antimycotic at 37°C for 4 hours. The resulting suspension was then poured through a 100µm cell strainer (Biologix, 15-1100) into a 50mL conical tube (Corning, 352098). The strainer was then rinsed with DMEM/F12 and the cells were centrifuged at 180g for 5 minutes. Cells were resuspended in DMEM (Life technologies, 11965-084) supplemented with 10% fetal bovine serum (FBS) (Cytiva, SH30071.03) and 5X antibiotic-antimycotic. Cells were maintained in medium containing 5X antibiotic-antimycotic for the first three passages, after which the concentration was reduced to 1X.

### Generation of mouse dermal fibroblasts stably expressing MCPyV TAs

To generate a lentiviral vector for expressing MCPyV tumor antigens, the truncated MCPyV TA sequence from the WaGa cell line was cloned into a modified version of the pTRIPZ lentiviral vector under control of the human EF1a promoter. A puromycin resistance gene under control of an internal ribosome entry site was placed downstream of the TA gene. To generate lentivirus, HEK293T cells at 70% confluence in a 10cm dish were transfected with 2.5µg the envelope plasmid pMD2.G, 7.5µg of the packaging plasmid pSPAX2, and 10µg of either the lentiviral vector containing MCPyV TAs or an empty vector control. Transfection was performed using lipofectamine 2000 (Thermo fisher scientific, 11668-027). Briefly, 45µL of lipofectamine 2000 was added to 500µL Opti-MEM (Life technologies, 31985-062) and incubated for 5 minutes at room temperature. Following the incubation, an additional 500µL of Opti-MEM containing the plasmid DNA was mixed into the solution. The transfection mixture was incubated for 20 minutes at room temperature before being added directly to the HEK293Ts. After 16-18 hours, the media on the HEK293Ts was replaced with fresh media. At 48 hours post-transfection, the supernatant was filtered through 0.45µm PVDF filters and hexadimethrine bromide (Sigma, H9268) was added to a final concentration of 6µg/mL. The supernatant was then added to MDFs in culture and incubated for 24 hours before the media was changed. Cells were selected with 4 µg/mL of puromycin (Sigma, P8833) for 1 week.

### Soft agar assay and clonal cell line isolation

Anchorage independent growth was assessed using a soft agar colony forming assay. The base layer for the soft agar consisted of 0.6% agarose (Seaplaque, 50100) in DMEM supplemented with 10% FBS and 100µg/mL Penicillin-Streptomycin (Life technologies, 15140-122) which was poured into plates at 50°C and left to solidify at room temperature for 30 minutes. Cells at about 70% confluence were washed in DPBS and detached by incubation in 0.05% Trypsin-EDTA (Life technologies, 25300-120) for 2 minutes at 37°C. Cells were then counted using a hemocytometer and diluted to an appropriate concentration in a solution of 0.4% agarose in DMEM supplemented with 10% FBS and 100µg/mL Penicillin-Streptomycin at 42°C. The cells were added on top of the base layer and cooled at room temperature for 30 minutes before the plates were transferred to a cell culture incubator at 37°C. Fresh media was added on top of the agarose layer every 2-3 days. To image colony formation, plates were stained with 0.005% crystal violet in PBS for 1 hour. To isolate clonal populations from the soft agar layer, colonies were gently pipetted out of the soft agar layer and placed into warm DMEM supplemented with 10% FBS. The colonies were observed in the plate to ensure that each well contained only a single colony. After colonies attached and cells grew to a confluent monolayer, cells were detached using 0.05% Trypsin-EDTA for 2 minutes at 37°C and expanded in DMEM supplemented with 10% FBS.

### Cell proliferation monitoring

Proliferation of some cell lines was monitored for extended periods of time. 10,000 cells were plated in one well of a 6-well plate in DMEM with 10% FBS. The cell culture medium was changed every other day and every sixth day the cells were washed with DPBS and detached by incubation in 0.05% Trypsin-EDTA for 2 minutes at 37°C. Cells were then counted using a hemocytometer, and 10,000 cells were plated into a new well of a 6-well plate in DMEM supplemented with 10% FBS.

### Western blotting

Cells for western blotting were lysed in a buffer with a pH of 7.5 containing 10mM HEPES, 500 mM NaCl, 1 mM EDTA, 1 mM DTT, 0.5% Triton X-100,1X PhosSTOP (Roche, 4906845001), and cOmplete protease inhibitors (Roche, 11836170001) by a 60-minute incubation at 4°C. Lysates were centrifuged for 10 minutes at 12,000g and 4°C to pellet insoluble material. Lysates were then mixed with Laemmli sample buffer, boiled for 2 minutes, and analyzed using SDS-PAGE. Western blot membranes were incubated with blocking buffer (5% milk in PBS with 0.1% Tween-20) for 1 hour at room temperature. Membranes were then incubated overnight at 4°C with primary antibodies diluted in blocking buffer. Membranes were washed once in PBS, twice in PBS with 0.1% Tween-20 and then incubated with HRP-linked secondary antibodies for 1 hour at room temperature. Membranes were washed once in PBS, twice in PBS with 0.1% Tween-20, and the signal was developed using the SuperSignal West Pico PLUS chemiluminescent substrate (Thermo fisher, 34580). Primary antibodies: anti-GAPDH (RRID:AB_561053, 1:5,000, CST 2118), anti-MCPyV TA (1:500, 2T2, Sigma MABF2316), anti-CK20 (RRID:AB_3086478, 1:10,000, Proteintech 82428-1-RR), anti-ATOH1 (RRID:AB_2861575, 1:1000, AbClonal A11477), anti-SOX2 (RRID:AB_358009, 1:500, R&D Systems MAB2018), anti-SYP (RRID:AB_2924257, 1:1000, CST 25056), anti-NSE(RRID:AB_2099180, 1:5000, Proteintech 10149-1-AP), anti-CHGA (RRID:AB_2081122, 1:1000, Proteintech 10529-1-AP), anti-NCAM1 (RRID:AB_2149421, 1:5000, Proteintech 14255-1-AP), anti-p53 (RRID:AB_331743, 1:1000, CST 2524), anti-pp53 (RRID:AB_331464, 1:1000, CST 9284), anti-human p21 (RRID:AB_823586, 1:1000, CST 2947), anti-mouse p21 (RRID:AB_628073, 1:200, SCBT sc-6246), anti-Rb (RRID:AB_632339, 1:1000, SCBT sc-50), anti-pRb (RRID:AB_11178658, 1:1000, CST 8516), anti-CCNB1 (RRID:AB_627338, 1:200, SCBT sc-245), anti-PCNA (RRID:AB_303062, 1:200, Abcam ab2426). anti-anti-Secondary antibodies: Anti-Mouse IgG, HRP-linked antibody (RRID:AB_330924, 1:2500, CST 7076), Anti-Rabbit IgG, HRP-linked antibody (RRID:AB_2099233, 1:2500, CST 7074).

### P53 activation assay

To activate p53, cells in phosphate buffered-saline were subjected to 1mJ/cm2 of UV-C in a Biorad GeneLinker GS chamber. Cells were then incubated in DMEM supplemented with 10% FBS (MDF cells) or RPMI supplemented with 20% FBS (MCC Cells) for 24 hours prior to collection for western blotting.

### Immunofluorescent staining of cells

Cells for immunofluorescent staining were seeded onto glass coverslips at a confluence of ∼50%. After 16 hours, the cells were washed in DPBS and fixed using 3% paraformaldehyde for 15 minutes at room temperature. Coverslips were incubated in blocking buffer (3% bovine serum albumin in PBS with 0.5% Triton X-100) for 1 hour at room temperature. The coverslips were then incubated with primary antibodies diluted in blocking buffer for 1 hour and washed three times with blocking buffer. Following the washes, coverslips were incubated with secondary antibodies diluted in blocking buffer for 1 hour and washed twice in blocking buffer. Coverslips were then incubated with DAPI in PBS for 5 minutes and then washed once in PBS. Coverslips were mounted onto glass slides using Fluoromount G mounting medium and dried overnight before imaging by fluorescence microscopy. Primary antibodies: anti-Vimentin (RRID:AB_10695459, 1:150, CST 5741) anti-MCPyV large tumor antigen (1:200, SCBT CM2B4), anti-Cytokeratin-20 (RRID:AB_3086478, 1:100, Proteintech 82428-1-RR), anti-SOX2 (RRID:AB_358009, 1:100, R&D systems MAB2018), anti-E-cadherin (RRID:AB_2291471, 1:100, CST 3195). Secondary antibodies: AlexaFluor 488 goat anti-mouse IgG H+L (RRID:AB_2534088, 1:500, Life technologies A11029), AlexaFluor 594 goat anti-rabbit IgG H+L (RRID:AB_2534079, 1:500, Life technologies A11012). Samples were imaged under conditions that produced minimal signal in samples that were stained only with secondary antibodies and no primary antibody (Figure S3A).

### RT-qPCR

Cells were washed in DPBS and detached from culture dishes by incubation in 0.05% Trypsin-EDTA for 2 minutes at 37°C. Cells were then washed in PBS and centrifuged for 5 minutes at 400g. RNA was extracted using the standard protocol for the Machery-Nagel Nucleospin RNA extraction kit and quantified using a NanoDrop spectrophotometer. cDNA synthesis was carried out using 17.5ng/µL RNA, 25ng/µL Oligo(dT) (Thermo fisher, SO132), 0.5mM dNTPs (Thermo fisher, 18427088), 1X MMLV reaction buffer (Promega, M531A), 10u/µL of MMLV reverse transcriptase (Promega, M170A), and 2u/µL RNAseOUT (Thermo fisher, 10777019). RNA was first incubated with dNTPs and Oligo(dT) for 5 minutes at 70°C before being cooled to 4°C for 2 minutes. After the reaction was cooled, the remaining reagents were added, and the reaction was incubated at 37°C for 50 minutes followed by a 15-minute incubation at 70°C. cDNA was cooled to 4°C and then aliquoted and frozen at -20°C. To measure gene expression, 0.4µL of cDNA was added to 1X PowerUP Sybr Green master mix (Thermo fisher, A25742) with 250nM of forward and reverse primers in a 20µL reaction volume and analyzed using a QuantStudio 3 instrument. qPCR reactions were done using the fast-cycling conditions described in the manufacturer’s instructions for the PowerUP Sybr Green master mix. All genes were measured in triplicate and normalized to GAPDH expression using the formula: GAPDH cycle threshold – Target Gene cycle threshold. All primers were synthesized by Integrated DNA Technologies (IDT). Previously published data concerning MCPyV tumor antigen transcript splicing was used to develop sT-specific primers(32).

### RNA-seq

Bulk RNA sequencing was performed using Plasmidsaurus. Cells were washed in PBS and then detached using a non-enzymatic detachment solution (Sigma, C5914). Cells were counted using a hemocytometer, and 100,000 cells were pelleted and resuspended in 50µL of RNA/DNA Shield (Zymo, R1100-50). The samples were sent to Plasmidsaurus where RNA was extracted and sequenced using a 3’ end counting approach with Illumina library prep including unique molecular identifiers (UMIs). Reads were filtered using FastP with poly-X tail trimming, 3’ quality-based tail trimming, a minimum Phred quality score of 15, and a minimum length requirement of 50 bp. Reads were then aligned to the reference genome using STAR aligner and BAM files were coordinate sorted using samtools. UMI based deduplication was done using UMICollapse. FASTQ files containing deduplicated reads were generated using samtools and transcripts were quantified using Salmon. Transcript quantification data was imported into RStudio using the tximeta package and summarized to gene-level data. The counts were then normalized using DESeq2’s vst function. Genes were ranked for fast gene set enrichment analysis (fgsea) by taking the difference in the normalized counts between the two groups being compared. The fgsea R package was used to determine if there was gene enrichment in MSigDB’s hallmark gene sets and graphs were generated using ggplot2.

### Mouse experiments

C57BL/6 (RRID:IMSR_JAX:000664) and NOD-scid-gamma mice (NSG) (RRID:IMSR_JAX:005557) were obtained from Jackson Labs. All animal work was approved by the University of Pennsylvania’s Institutional Animal Care and Use Committee. All cell lines used in mice were tested and verified to be pathogen free by IDEXX. To generate tumors in mice, cells were washed in DPBS and detached using 0.05% Trypsin-EDTA for 2 minutes at 37°C. Cells were counted using a hemocytometer and viability was confirmed to be >90% using trypan blue staining. Cells were washed twice in DPBS, resuspended in DPBS, and cooled on ice. An equal volume of ice-cold growth-factor reduced Matrigel (Corning 354230) was added to the cells. 100µL containing either 1x10^6 or 2x10^6 cells in the DPBS-matrigel mixture were subcutaneously injected into the right flank of the mice. Mice were monitored at least every other day following tumor cell injections. Tumors were measured using digital calipers and the volume was calculated using the formula 0.5 x Length x Width². Mice were euthanized when they had a body condition score of 1, tumor volume of 2cm³, any tumor dimension reached 2cm, impaired mobility due to tumor growth, exhibited hunched posture with ruffled fur, or exhibited significant lethargy. Tumors are named after the cell line used to generate the tumor, and subsequent tumor cells isolated from each tumor are considered the next generation of the tumor. The MDF-TA-12 cells isolated from soft agar are considered the F0 cell line, which is used to generate F0 tumors. The F1 cell line was isolated from an F0 tumor and used to generate the F1 tumors. The F2 cell line was isolated from an F1 tumor and used to generate F2 tumors. The F3 cell line was isolated from an F2 tumor.

### Tumor tissue processing

Tumor tissue was fixed in 10% neutral-buffered formalin for 24 hours. The tissue was then transferred to 70% ethanol and embedding and sectioning were carried out by the Wistar Histotechnology Core. Slides were deparaffinized by two 10-minute incubations in Xylene, followed by two 5-minute incubations in absolute ethanol, one 5-minute incubation in 80% ethanol, one 5-minute incubation in 50% ethanol, one 5-minute incubation in 30% ethanol, and two 5-minute incubations in water. Rehydrated tumor tissue was stained with hematoxylin and eosin (Abcam AB245880) according to the manufacturer’s instructions.

### Immunofluorescent staining of tumor tissue

To probe for MCC markers in tumor sections, rehydrated tumor tissue was subjected to heat-induced epitope retrieval by incubation in a boiling antigen retrieval solution consisting of 10mM Tris, 1mM EDTA, 0.1% Triton X-100 at pH 9. Samples were boiled for 25 minutes followed by cooling to room temperature over 1 hour. After the samples had cooled, they were washed in PBS and then blocked using 3% bovine serum albumin in PBS with 0.5% Triton X-100 for 1 hour. Samples were then incubated with primary antibody for 1 hour before being washed three times in PBS. Samples were then incubated with secondary antibody and DAPI for 1 hour before being washed three times in PBS. Samples were then briefly dipped in distilled water and Flouromount G aqueous mounting medium was added to the tissue sections. A glass coverslip was placed on the sections, and they were left in the dark overnight to dry. Images were acquired using fluorescence microscopy and all images were acquired under conditions that produced minimal signal in tissue stained only with secondary antibodies. Primary antibodies: anti-MCPyV large tumor antigen (1:200, SCBT CM2B4), anti-Cytokeratin-20 (RRID:AB_3086478, 1:100, Proteintech 82428-1-RR). Secondary antibodies: AlexaFluor 488 goat anti-mouse IgG H+L (RRID:AB_2534088, 1:500, Life technologies A11029), AlexaFluor 594 goat anti-rabbit IgG H+L (RRID:AB_2534079, 1:500, Life technologies A11012). Samples were imaged under conditions that produced minimal signal in samples that were stained only with secondary antibodies and no primary antibody (Figure S3B).

### Tumor cell isolation

Tumor tissue for cell isolation was collected into DMEM with 10% FBS and 5X antibiotic-antimycotic. The tissue was washed in DPBS and the cells were isolated in the same manner used to isolate mouse dermal fibroblasts from mouse ear tissue.

### Image analysis and quantification

Tissue sections were analyzed using TissueLab(33), an integrated medical imaging analysis platform supporting deep learning-based and custom computational pipelines for high-throughput tissue quantification, to assess expression of the MCC tumor marker CK20 across multiple sections.

The analysis proceeded in three primary stages: segmentation, spatial masking, and intensity quantification. Nuclei segmentation was performed on DAPI-stained sections using a deep learning-based approach, generating precise contour coordinates for each detected nucleus, including those in high-density cellular regions. Quantification was carried out using a custom Python-based analysis pipeline integrated within the TissueLab platform.

For the cytoplasmic marker CK20, which exhibits both cytoplasmic and perinuclear localization, a morphological dilation strategy was employed to detect and quantify the perinuclear CK20 signal. Specifically, a 15-pixel perinuclear ring was generated by dilating each nuclear contour and subtracting the original nuclear area, yielding a standardized region for measuring average CK20 pixel intensity.

Tumor marker positivity for individual cells was determined using fixed intensity thresholds (CK20 ≥ 15), established based on global intensity distributions across the dataset. These thresholds were applied uniformly across all samples to enable consistent comparative analysis of tumor-positive cell populations.

## Results

### MCPyV tumor antigens induce oncogenic transformation of primary dermal fibroblasts

To determine whether MCPyV TAs can induce oncogenic transformation in primary dermal fibroblasts, MDFs were engineered to express MCPyV T antigens using a lentiviral vector containing the truncated T antigen sequence from the MCC cell line WaGa. This specific T antigen sequence was chosen because WaGa is considered to be representative of MCC tumors(34). Stable TA-expressing cells (MDF-TA) were generated by puromycin selection (Figure 1A). Expression of the TAs lead to increased growth rate in MDF-TA cells relative to the vector control cells (MDF-empty) (Figure 1B). The starting MDF population and the transduced cells were positive for expression of the dermal fibroblast marker vimentin and negative for the epithelial marker e-cadherin, consistent with a fibroblast-phenotype but different from the epithelial WaGa cells (Figure 1C, S1A, S1B). The MDF-TA cells exhibited an altered morphology compared with vector-only control cells (Figure 1C). To assess oncogenic transformation, the MDF-TA cells were evaluated for anchorage-independent growth in soft agar. MDF-TA cells formed colonies in soft agar, whereas control cells did not, indicating the acquisition of a transformed phenotype (Figure 1D). Clonal cell lines were established by isolating individual colonies from soft agar cultures (Figure 1E). Two representative clones, MDF-TA-4.1 and MDF-TA-12, exhibited increased growth and altered morphology relative to the parental MDF-TA population (Figure 1B, 1C). The isolated cells also exhibited faster growth in soft agar and formed clearly visible colonies after 21 days, whereas the parental MDF-TA cells produced few detectable colonies under the same conditions (Figure 1F). Consistent with the increased growth rate of MDF-TA-12, MDF-TA-12 colonies were much larger than those of MDF-TA and MDF-TA-4.1(Figure 1G). Anchorage-independent growth, along with an increase in growth rate, demonstrates that expression of MCPyV tumor antigens is sufficient to induce oncogenic transformation of primary MDFs.

**Figure 1.**
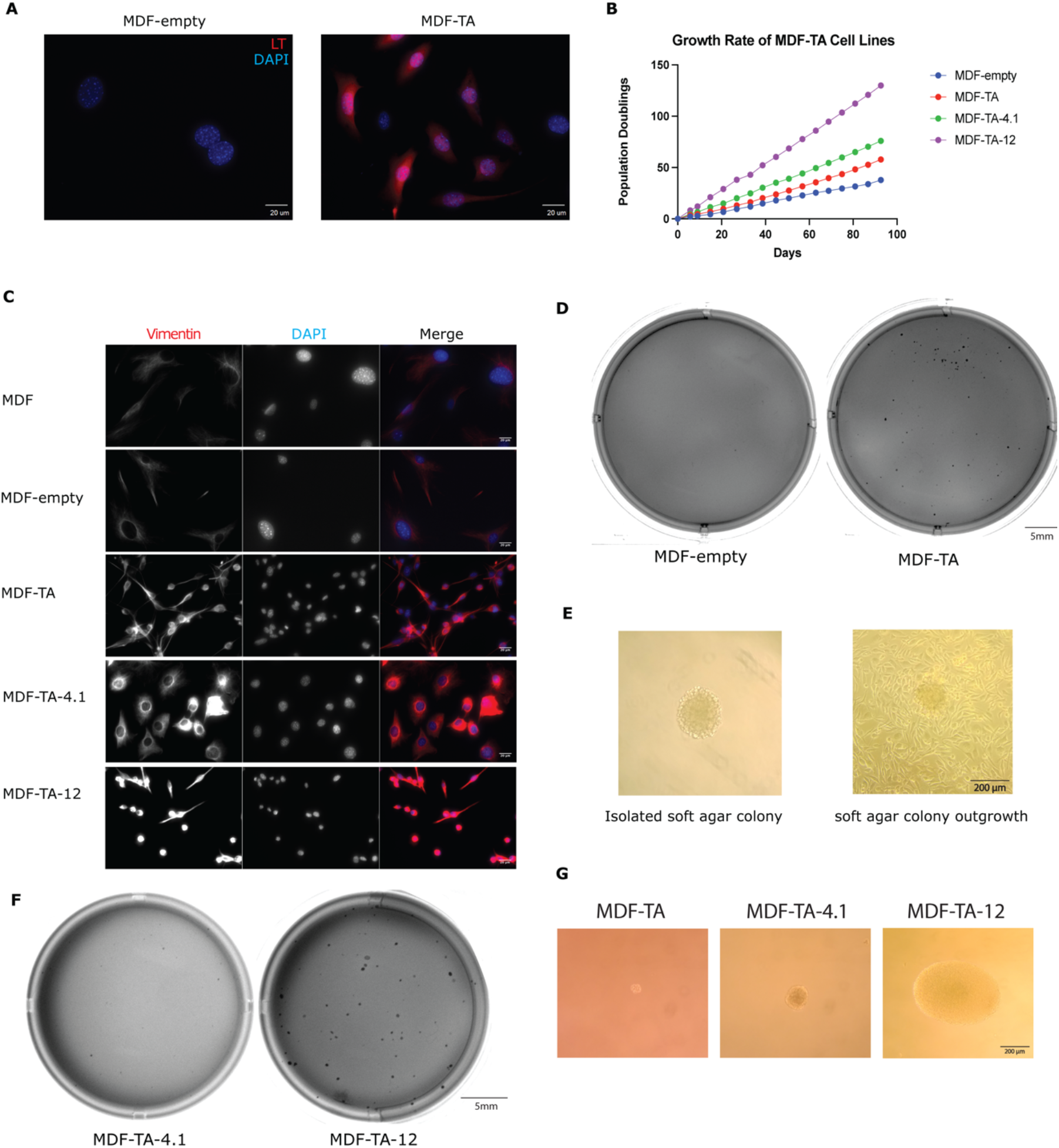
MCPyV TAs transform mouse dermal fibroblasts. (A) Immunofluorescent staining of MCPyV LT in MDFs transduced with a vector control (MDF-empty) or MCPyV T antigens (MDF-TA). Images were taken at 40X magnification. (B) Population doublings were tracked over a period of 3 months in MDF-empty cells, MDF-TA cells, MDF-TA-4.1 cells, and MDF-TA-12 cells. (C) Immunofluorescent staining of vimentin shows the morphology of MDFs, MDF-empty cells, MDF-TA cells, MDF-TA-4.1 cells, and MDF-TA-12 cells. Images were taken at 40X magnification. (D) Crystal violet staining of soft agar plates containing 2500 MDF-TA and MDF-empty cells after 30 days of growth in 6-well plates, colony formation is visible in MDF-TA cells and not observed in MDF-empty cells. (E) (left) Representative image of an isolated soft agar colony after removal from the soft agar plate and (right) representative image of outgrowth after 4 days in culture. (F) Crystal violet staining of soft agar plates containing 1000 MDF-TA-4.1 and MDF-TA-12 after 21 days of growth in 12-well plates, showing variability in the capacity of isolated clonal cell lines to grow in soft agar conditions. Images are representative of triplicate wells. (G) Representative images of soft agar colonies formed by MDF-TA, MDF-TA-4.1, and MDF-TA-12 cells after 19 days of growth in soft agar.

**Table 1.**
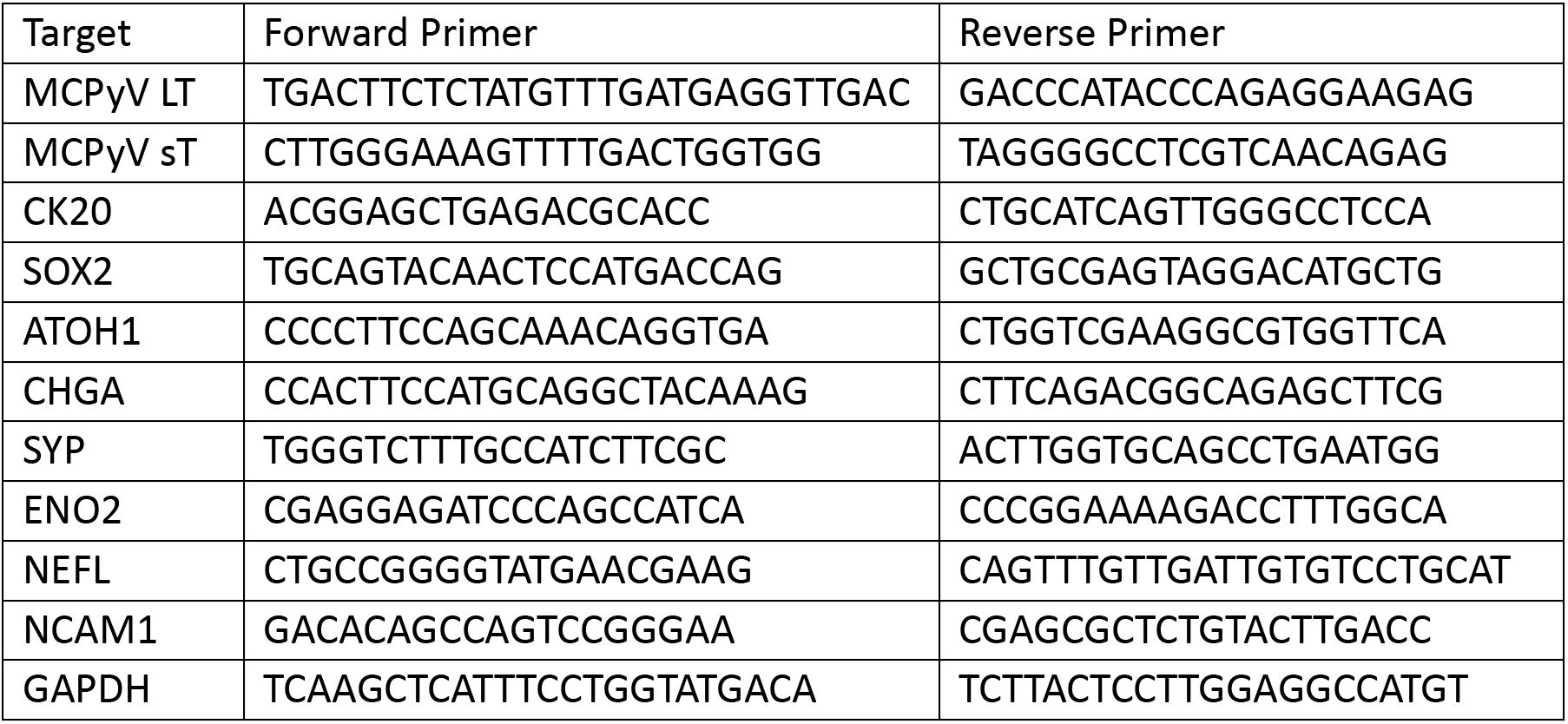
Primer Sequences.

### Mouse dermal fibroblasts expressing T antigens acquire molecular features of MCC

After observing that TA expression in MDFs induced oncogenic transformation, we next asked whether these cells had also acquired molecular features characteristic of MCC. Western blot analysis showed much higher LTT expression in all TA-expressing MDF cell lines (MDF-TA, MDF-TA-4.1, and MDF-TA-12) compared to the MCC cell lines, WaGa and MKL-1, whereas sT protein was not detected in either the transformed cell lines or the MCC cell lines (Figure 2A). We next examined expression of canonical MCC markers. CK20 protein was weakly detected in the parental MDF-TA cells and strongly expressed in the MDF-TA-12 clone (Figure 2A). SOX2 protein was detected in all TA-expressing MDF cell lines but was undetectable in the vector control cells (Figure 2A). In contrast, expression of Merkel cell lineage regulator atonal homolog 1 (ATOH1) protein was not detected in any of the murine cell lines (Figure 2A). ATOH1 mRNA was likewise undetected by RT-qPCR. At the transcript level, TA-expressing MDFs exhibited increased expression of neuroendocrine-associated genes, including KRT20, SOX2, CHGA, NCAM1, ENO2, SYP, and NEFL, although the degree of expression varied among the individual TA-expressing cell lines and low levels of SOX2, NCAM1, and ENO2 were detected in the vector control cells (Figure 2B-H). Despite detectable mRNA, only SOX2, CK20, and ENO2 were readily detectable at the protein level (Figure 2A, 2D-2H). Immunofluorescent staining was used to confirm expression of SOX2 and CK20. SOX2 was detected in all TA-expressing MDF cell lines, whereas CK20 expression was heterogenous, with the strongest signal observed in the MDF-TA-12 clone (Figure 2I, S2A-B, S3A). These findings demonstrate that TA expression in MDFs can induce multiple molecular features characteristic of MCC, although not all canonical MCC markers are uniformly induced.

**Figure 2.**
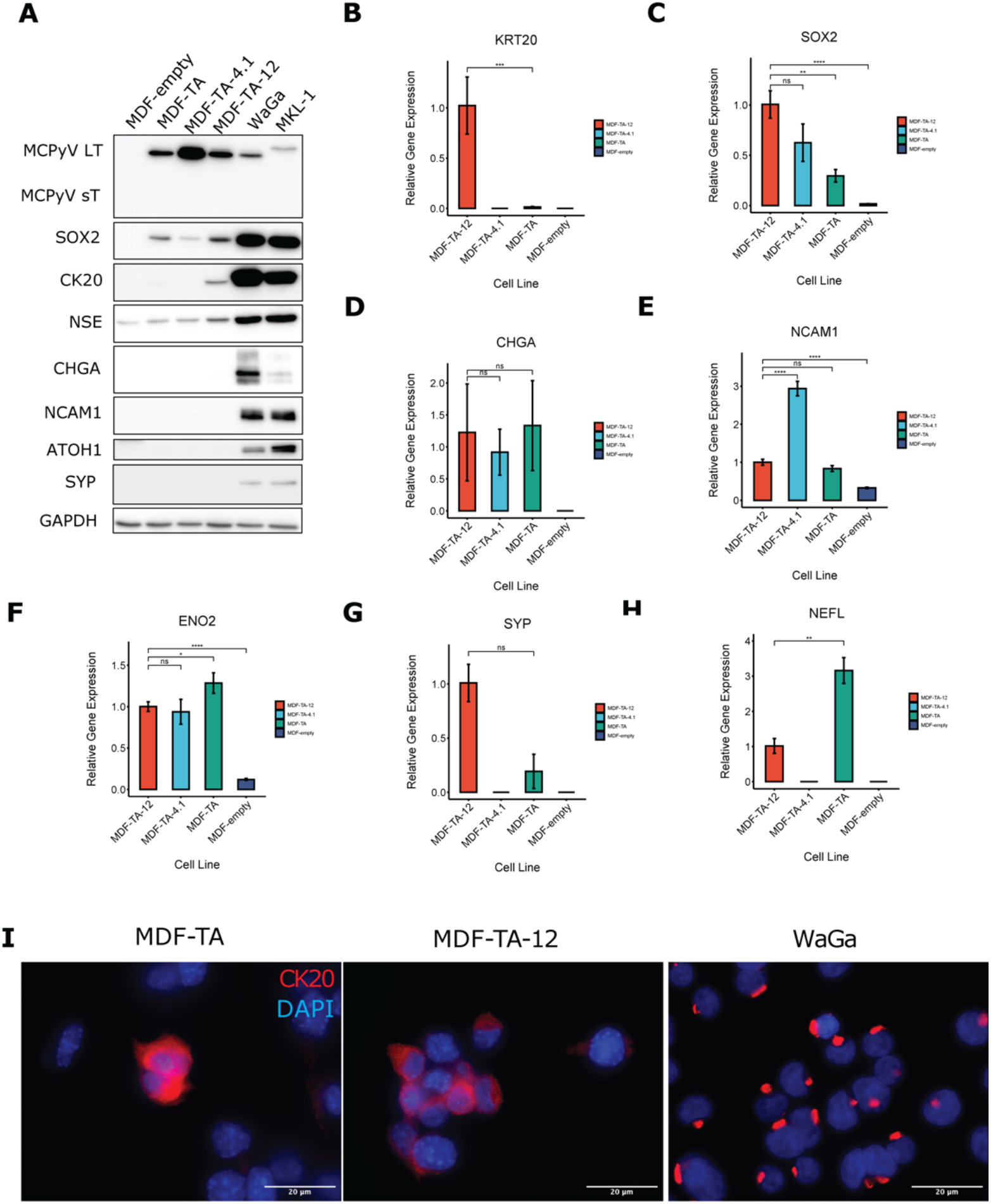
MCPyV tumor antigens induce expression of MCC lineage markers. (A) Expression of the MCC markers large tumor antigen (LT), small tumor antigen (sT) cytokeratin 20 (CK20), sry-box transcription factor 2 (SOX2), neuron-specific enolase 2 (NSE), chromogranin A (CHGA), neural cell adhesion molecule 1 (NCAM1), atonal homolog 1 (ATOH1), and synaptophysin (SYP) in MDF cell lines was assessed by western blot. SOX2 expression was detected in all TA-expressing cells, while CK20 was weakly detected in MDF-TA and detected in MDF-TA-12 cells. Lysates from two MCC cell lines, WaGa and MKL-1, were included as positive controls for MCC marker expression. GAPDH served as a loading control. (B-H) mRNA Expression of KRT20, SOX2, CHGA, NCAM1, ATOH1, CHGA, ENO2, SYP, and NEFL measured by RT-qPCR. All RT-qPCR data were normalized by subtracting the target gene Ct value from the Ct value for GAPDH to get the ΔCt value, and gene expression relative to MDF-TA-12 was calculated using the formula 2^-ΔΔCt^. Samples with at least 2 out of 3 replicates having Ct values >35 were considered to be undetected. Error bars represent the standard deviation. P values were calculated by comparing the ΔCt values between samples using Welch’s t test. Statistical comparisons were not performed for undetected samples. Significant differences are indicated as follows: * indicates p < 0.05, ** indicates p<0.01, *** indicates p<0.001, **** indicates p<0.0001. (I) Immunofluorescent staining of MDF-TA, MDF-TA-12, and WaGa cells for the MCC marker CK20. Images acquired under the same conditions at 100X magnification.

### TA-expressing MDFs exhibit transcriptional programs associated with MCC

To characterize transcriptional changes associated with MCPyV T antigen expression, we performed bulk RNA-sequencing of MDF-TA, MDF-TA-4.1, MDF-TA-12, and control MDF-empty cells. Gene set enrichment analysis using the MSigDB hallmark gene sets revealed strong enrichment of G2/M checkpoint and E2F transcriptional programs in MDF-TA cells compared with controls (Figure 3A). MDF-TA-4.1 and MDF-TA-12 cells both showed enrichment in E2F transcriptional activity relative to the parental MDF-TA cells (Figure S4A, S4B). The parental MDF-TA population displayed enrichment of interferon-related gene expression, consistent with previous reports that MCPyV T antigen expression can induce inflammatory transcriptional responses(35) (Figure 3A). Interestingly, this phenotype was retained in only one clone, MDF-TA-4.1 (Figure 3A). Both soft agar-derived clones, MDF-TA-4.1 and MDF-TA-12, exhibited decreased expression of genes associated with epithelial-mesenchymal transition (EMT), indicating a shift towards a more epithelial phenotype (Figure 3A). Both isolated cell lines showed enrichment of MYC target genes, and MDF-TA-12 also displayed increased mTORC1 signaling gene set enrichment, consistent with MYC and mTOR pathway activation frequently observed in MCC(18, 36). Despite differences in morphology and growth rate, the two soft agar-derived MDF-TA cell lines only differed significantly in interferon-related gene expression and mTOR pathway activity (Figure S4C). Overall, the transcriptomic data indicate that TA expression in dermal fibroblasts can induce transcriptional programs associated with cell-cycle activation, inflammatory signaling, and oncogenic MYC/mTOR pathways.

**Figure 3.**
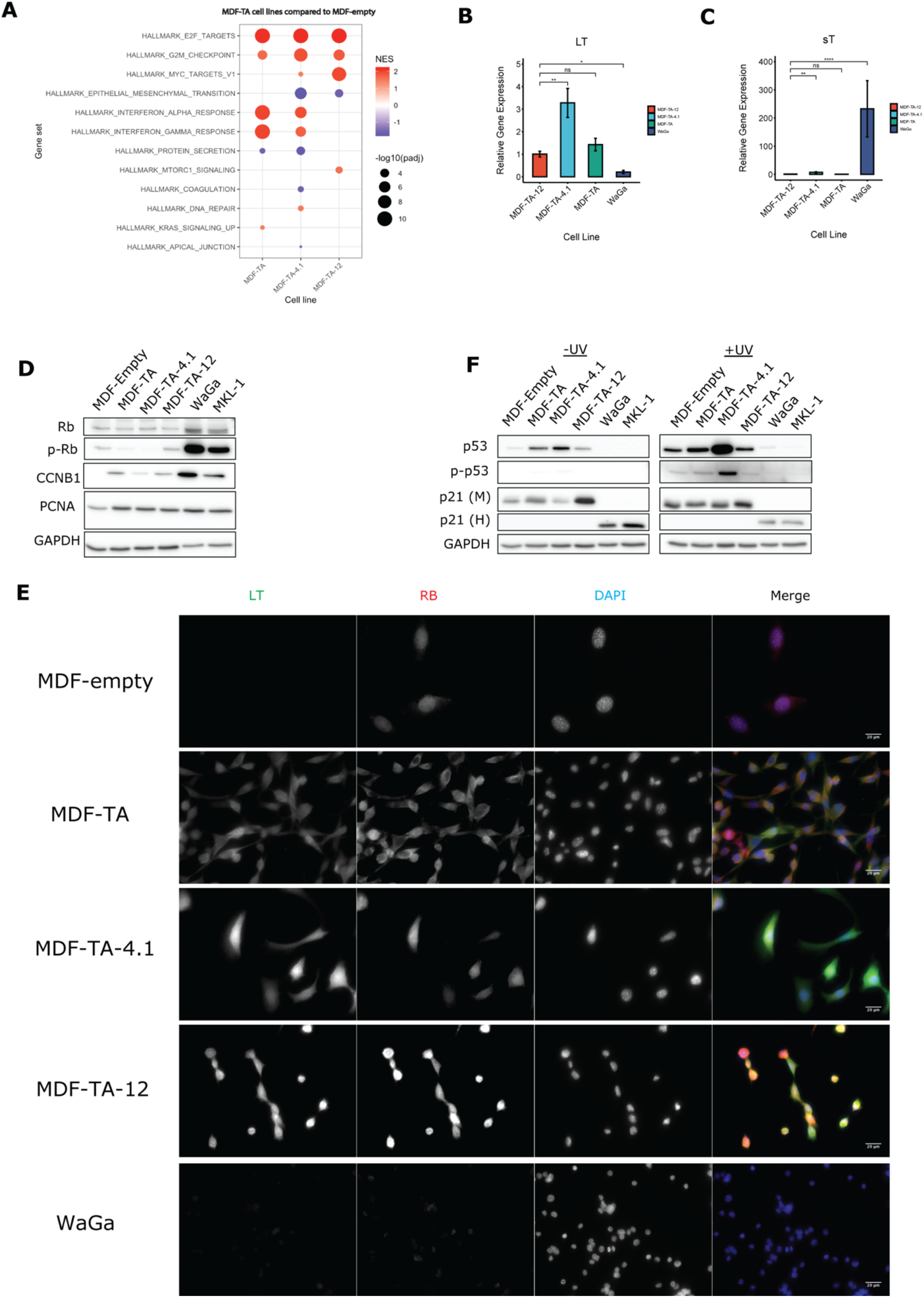
MDF-TA cells exhibit MCC-like transcriptional programs and Rb inhibition. (A) Fast gene set enrichment analysis (fgsea) of RNA-seq data from MDF-TA cell lines using the MSigDB Hallmark gene set collection. The net enrichment score (NES) of each gene set is represented by color, with red indicating positive enrichment and blue indicating negative enrichment. Dot size represents the –log10 value of the adjusted p-value (padj), such that larger dots indicate more statistically significant enrichment. (B, C) mRNA Expression of LT and sT measured by RT-qPCR. All RT-qPCR data were normalized by subtracting the target gene Ct value from the Ct value for GAPDH to get the ΔCt value, and gene expression relative to MDF-TA-12 was calculated using the formula 2^-ΔΔCt^. Samples with at least 2 out of 3 Ct values >35 were considered to be undetected. Error bars represent the standard deviation. P values were calculated by comparing the ΔCt values between samples using Welch’s t test. Significant differences are indicated as follows: * indicates p < 0.05, ** indicates p<0.01, *** indicates p<0.001, **** indicates p<0.0001. (D) Expression of Rb, p-Rb, CCNB1, and PCNA were assessed by western blot in MDF-empty, MDF-TA, MDF-TA-4.1, and MDF-TA-12. WaGa and MKL-1 were included as positive controls. (E) Immunofluorescent staining of LT and Rb in MDF cell lines and WaGa cells. Images acquired under identical conditions at 40X magnification. (F) Expression of p53, p-p53, and p21 were detected in MDF cells lines and MCC cell lines with and without exposure to UV-C radiation. p21(M) indicates mouse p21, while p21(H) indicates human p21. Blots containing UV-C treated and untreated samples were developed simultaneously.

### MCPyV T antigen expression inhibits Rb but not p53 in mouse dermal fibroblasts

LTT has previously been shown to inhibit the tumor suppressor Rb by binding via its LxCxE motif, while sT is thought to be responsible for inhibition of another key tumor suppressor, p53(15, 19). To determine whether the T antigens were exhibiting these functions in the MDFs, we first compared T antigen mRNA expression levels between MDF cell lines and WaGa cells using RT-qPCR. All TA-expressing MDF cell lines exhibited significantly increased levels of LTT but markedly decreased levels of sT when compared to WaGa cells (Figure 3B, 3C). Because LTT-mediated inhibition of Rb is expected to activate E2F-dependent transcription, we next examined expression of the E2F targets CCNB1 and PCNA(16). Both proteins were increased in all TA-expressing cells compared with the vector control cells, despite similar or lower levels of inhibitory Rb phosphorylation (Figure 3D). These findings are consistent with LTT-mediated inhibition of Rb occurring independently of Rb phosphorylation. Immunofluorescent staining showed colocalization of LTT and Rb in TA-expressing MDFs, consistent with the established mechanism where MCPyV LTT binds Rb to inhibit its function (Figure 3E).

To determine whether p53 remained functional in the MDF cells, we treated cells with UV-C radiation to induce DNA damage and activate p53. Basal p53 expression was elevated in TA-expressing MDFs relative to the vector control cells and increased further after UV-C treatment, accompanied by increased p53 phosphorylation (Figure 3F). These findings are consistent with previous reports that MCPyV LTT activates p53 signaling(19). In contrast, p53 was undetected in MCC cell lines prior to UV-C treatment, and only low levels of p53 were detected following UV-C exposure (Figure 3F). Expression of p21, a major transcriptional target of p53, was increased after UV-C treatment in MDF-empty, MDF-TA, and MDF-TA-4.1 cells, but not in MDF-TA-12, WaGa, or MKL-1 cells (Figure 3F). The absence of p21 induction in WaGa and MKL-1 cells is consistent with previous work demonstrating that p53 signaling is functionally suppressed in MCPyV-positive MCC through MDM2/MDM4-dependent mechanisms(19). The lack of p21 induction in MDF-TA-12 cells was accompanied by relatively high basal p21 expression, suggesting that p53 may already be active prior to UV-C treatment (Figure 3F). Consistent with the low level of sT expression observed in the TA-expressing MDFs (Figure 3C), sT expression may be insufficient to suppress p53 signaling in these cells. Together, these data demonstrate that TA-expressing MDFs recapitulate LT-mediated inhibition of the Rb pathway while retaining functional p53 signaling, suggesting that transformation of these cells is driven primarily by LT rather than sT.

### TA-transformed fibroblasts generate MCC-like tumors *in vivo*

To test whether TA-transformed fibroblasts were tumorigenic *in vivo*, MDF-TA-4.1 and MDF-TA-12 cells were injected subcutaneously into C57BL/6 mice. Injection of MDF-TA-12 cells resulted in palpable tumors in 5 of 6 mice by day 6 (Figure 4A). On day 6 and 7, three mice were euthanized and the tumors were removed to isolate tumor-derived cells (Figure 4B). In the remaining mice with MDF-TA-12 tumors, tumor growth peaked around day 8–9 but subsequently regressed, and viable tumor cells could no longer be recovered by day 12. Histological analysis demonstrated highly cellular tumors composed of poorly differentiated cells, consistent with human MCC morphology (Figure 4C).

**Figure 4.**
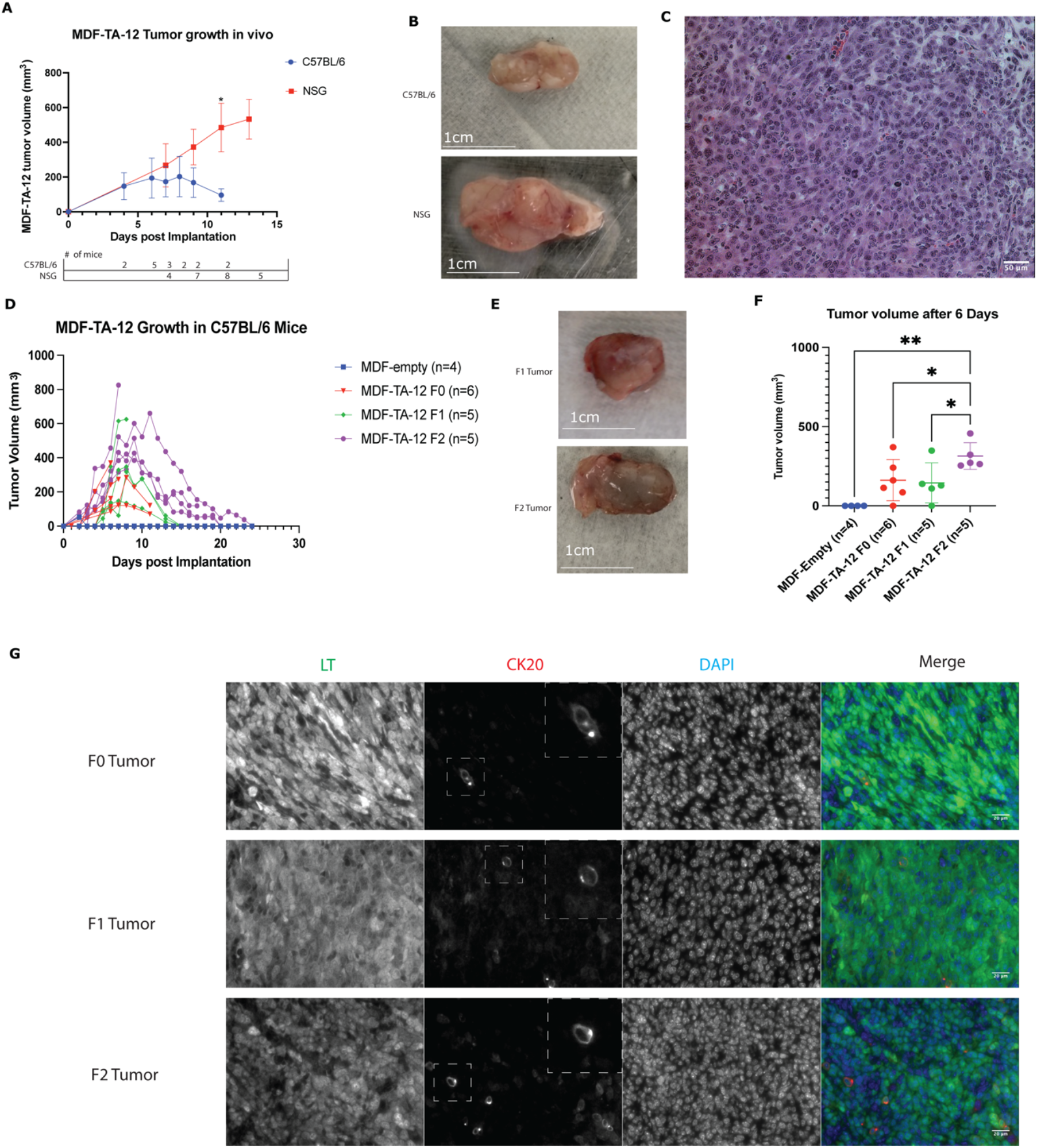
MDF cells transformed with MCPyV tumor antigens form MCC-like tumors in mice. (A) Growth of MDF-TA-12 tumors measured over time following implantation in NSG and C57BL/6 mice. Two C57BL/6 mice were euthanized on Day 6, one on Day 7, and two on Day 12. Three NSG mice were euthanized on Day 11, and five on Day 13. This graph includes data for mice that developed measurable tumors. The number of mice with tumors measured at any timepoint is indicated in the table directly below the chart. P values were calculated using the Mann Whitney test to compare C57BL/6 and NSG tumor sizes at each time point, * indicates P<0.05. B) Representative images of MDF-TA-12 tumors. These specific tumors were removed from a C57BL/6 mouse at day 7(top) and an NSG mouse at day 13 (bottom). (C) Hematoxylin and eosin (H&E) staining of a tumor isolated from a C57BL/6 mouse implanted with MDF-TA-12 cells. Image acquired at 20X magnification. (D) Growth curves of all tumors formed in C57BL/6 mice by MDF-empty, MDF-TA-12 F0, MDF-TA-12 F1, and MDF-TA-12 F2. (E) Images of an MDF-TA-12 F1 tumor removed from a C57BL/6 mouse on day 8 (top) and an MDF-TA-12 F2 tumor removed from a C57BL/6 mouse on day 7 (bottom). (F) Tumor volume for MDF-TA-12 F0, F1, and F2 for tumors measured 6 days post implantation. Tumor volume was calculated using the formula 0.5 x Length x Width². P values were calculated using Welch’s t test; * indicates P<0.05, ** indicates P<0.01. (G) Immunofluorescent staining of formalin-fixed, paraffin-embedded tumor tissue from the F0, F1, and F2 generation of tumors. Tumor tissue is stained for LT and CK20, images were acquired under identical conditions at 40X magnification. White dashed lines indicate selected areas of the CK20 staining are enlarged to highlight the perinuclear dot-like staining pattern, with the enlarged image in the upper-right hand corner of the CK20 image.

To determine whether tumor regression was due to immune-mediated clearance, MDF-TA-12 cells were injected into immunodeficient NOD-scid-gamma (NSG) mice. In this setting, palpable tumors were detected in 4 of 9 mice by day 7 and 8 of 9 mice by day 11 (Figure 4A, 4B). The tumors grew aggressively, with several mice reaching humane endpoints by day 11–13 (Figure S5A). In at least one mouse we observed invasion of surrounding muscle tissue (Figure S5B) and development of ascites containing tumor cells. Despite their aggressive growth in NSG mice, the MDF-TA-12-A cells (recovered from the ascites fluid of an NSG mouse) grew poorly when injected into C57BL/6 mice (Figure S5C). The ability of MDF-TA-12 cells to persist only in NSG mice suggests that immune surveillance plays a critical role in controlling growth of TA-transformed fibroblasts in immunocompetent hosts.

Because tumors generated by the parental MDF-TA-12 cells (F0) regressed in immunocompetent mice, we next asked whether serial *in vivo* passaging under immune pressure could select for tumor cells with enhanced fitness. Cells derived from the largest F0 tumor were reinjected into C57BL/6 mice as MDF-TA-12 F1 cells. Tumors formed in 4 of 5 mice but again regressed over time, with one mouse developing a tumor that reached a volume of over 600 mm³ before growth began to slow (Figure 4D).

The mouse from the second round of *in vivo* passage whose tumor exhibited the most aggressive growth was sacrificed at day 8. The F1 tumor was removed (Figure 4E) and used to generate MDF-TA-12 F2 cells. These cells formed tumors in all injected mice (Figure 4D) and produced significantly larger tumors by day 6 compared with earlier passages (Figure 4E-4F). MDF-TA-12 F2 tumors that were followed until complete regression persisted for an average of 21.3 days, compared with an average of 15 days for MDF-TA-12 F1 tumors. Although F2 tumors ultimately regressed in C57BL/6 mice (Figure 4D), these results demonstrate that serial *in vivo* passaging enhanced tumor growth and persistence in immunocompetent mice. Immunofluorescent staining of tumor tissue from each generation confirmed persistent LTT and CK20 expression throughout serial passaging (Figure 4G). Some tumor cells displayed the perinuclear dot-like CK20 pattern observed in human MCC, although this feature remained heterogeneous among tumors (Figure 4G). To quantify this heterogeneity, perinuclear CK20 staining was analyzed in three independent F0 tumors using TissueLab. CK20 positive cells ranged from <1% in the least CK20-positive tumor to 4.2% in the most CK20-positive tumor (Figure S6A-D), demonstrating intertumoral variability in CK20 expression. Together, these results establish that TA expression is sufficient to generate MCC-like tumors, which remain under strong immune control, providing a useful system for studying tumor evolution during immune selection.

### Serial *in vivo* passaging reveals molecular adaptations associated with tumor evolution under immune pressure

Having established that serial *in vivo* passaging enhanced tumor growth, we next asked whether immune selection was accompanied by molecular adaptations in the transformed cells. Cells isolated from the F2 generation of tumors were expanded to generate the F3 cell line. Compared with the parental F0 cells, F3 cells exhibited a significantly decreased expression of LTT together with a striking increase in sT expression, suggesting that there is selective pressure *in vivo* that favors expression of sT over LTT (Figure 5A, 5B). We next examined whether this transition was accompanied by changes in pathways involved in immune surveillance.

**Figure 5.**
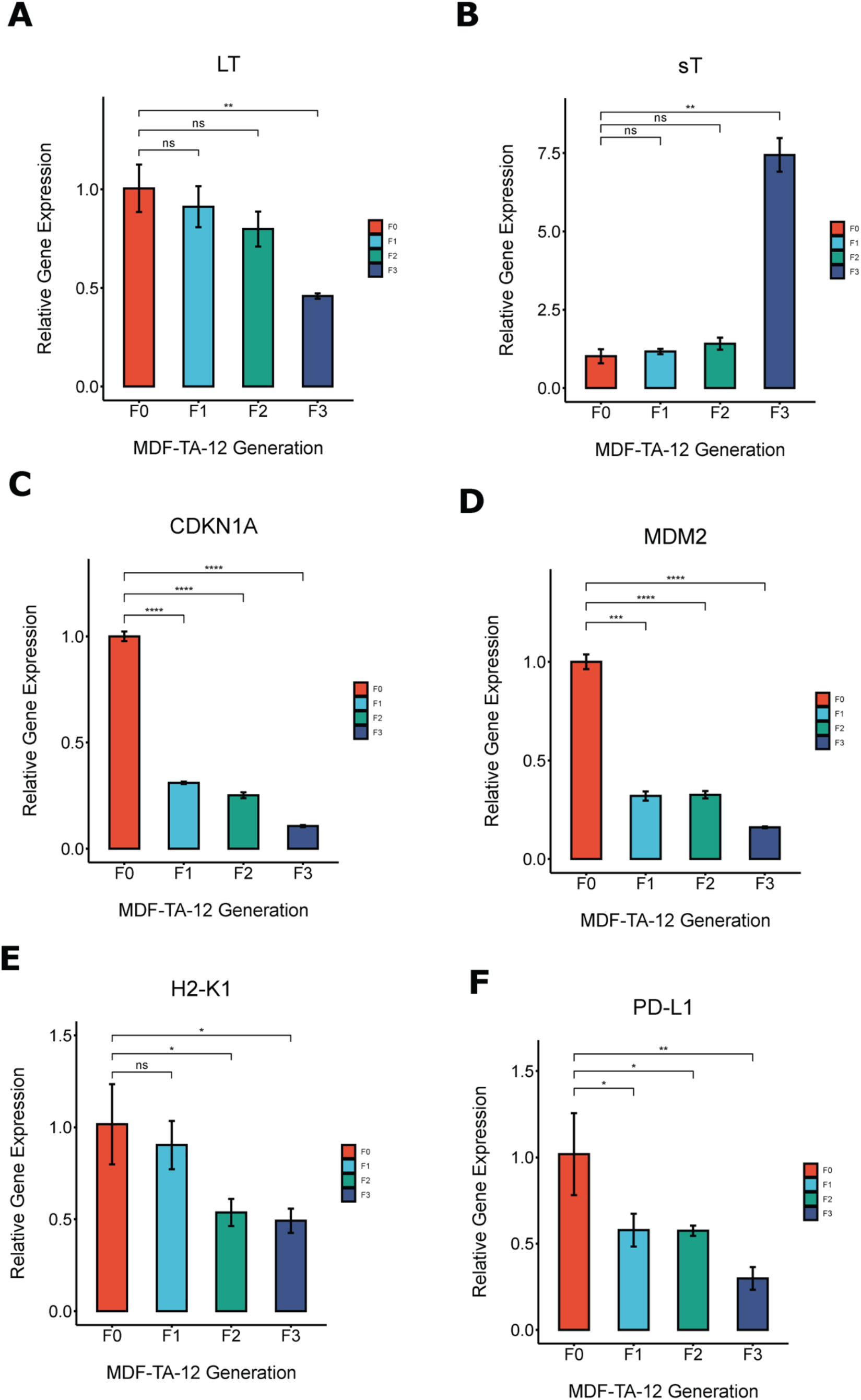
Transcriptional changes acquired by MDF-TA-12 cells after serial *in vivo* passage. (A-F) mRNA Expression of LT, sT, CDKN1A, MDM2, H2-K1, and PD-L1 measured by RT-qPCR. All RT-qPCR data were normalized by subtracting the target gene Ct value from the Ct value for GAPDH to get the ΔCt value, and gene expression relative to MDF-TA-12 F0 was calculated using the formula 2^-ΔΔCt^. Samples with at least 2 out of 3 Ct values >35 were considered to be undetected. P values were calculated by comparing the ΔCt values between samples using Welch’s t test. Significant differences are indicated as follows: * indicates p < 0.05, ** indicates p<0.01, *** indicates p<0.001, **** indicates p<0.0001.

Expression of the p53 target genes CDKN1A and MDM2 was significantly decreased following the first round of *in vivo* passaging and declined further in the F3 generation of cells concurrent with the increase in sT expression (Figure 5C, 5D). Similarly, expression of the murine MHC-1 gene H2-K1 progressively decreased during serial passaging, while PD-L1 expression also declined relative to the F0 generation (Figure 5E, 5F). Overall, these findings demonstrate that prolonged growth under host immune pressure is accompanied by coordinated transcriptional changes involving T antigen isoform expression, p53 signaling, and antigen presentation pathways. Importantly, the transcriptional changes arising in the cells after *in vivo* passaging highlight key adaptations that MCPyV-driven tumors can acquire in response to immune pressure.

## Discussion

MCPyV tumor antigens are essential drivers of MCPyV+ MCC, but many aspects of MCPyV - driven oncogenesis remain unclear. MCPyV+ tumors don’t usually possess obvious oncogenic mutations outside of MCPyV integration, yet transformation of primary skin cells by MCPyV TAs alone has not been demonstrated(9, 10). Our data demonstrate that expression of the MCPyV TAs is sufficient to transform primary MDFs into cells that share molecular and phenotypic similarities with MCC. The TA-transformed MDFs exhibited anchorage-independent growth, expressed MCC-associated markers, displayed transcriptomic changes characteristic of MCC, and formed tumors *in vivo*. Unlike most previously described models, which rely on epithelial lineage reprogramming or additional engineered oncogenic alterations to achieve tumor formation, expression of MCPyV T antigens alone was sufficient to generate MCC-like tumors. This enabled us to monitor tumor progression without introducing additional genetic drivers. Our work has thus established an immunocompetent model for studying MCPyV-driven tumorigenesis and tumor evolution under immune pressure.

We found that the transformed fibroblasts acquired multiple MCC-like molecular features. Notably, we detected expression of the key MCC markers CK20 and SOX2 (Figure 2A, 2B, 2C). Expression of the neuroendocrine genes CHGA, ENO2, NEFL, SYP, and NCAM1 was detected in the transformed cells as well, although expression was generally low and could not be confirmed at the protein level for all neuroendocrine genes (Figure 2A, 2D-2H). In contrast, these cells lacked expression of ATOH1, the classical Merkel cell lineage transcription factor (Figure 2A). Although ATOH1 is considered a key regulator of the Merkel cell lineage, Fan et al. demonstrated that ATOH1 is not actually required for proliferation of MCC cells(37). Moreover, ATOH1 expression varies substantially among MCC tumors, and higher ATOH1 expression has been associated with more advanced disease (38). Despite lacking ATOH1, this model recapitulates many key features of MCPyV-positive MCC while providing a genetically simple, immunocompetent system for studying MCPyV-driven tumor evolution under host immune selection. We detected expression of both LTT and sT in the transformed cells, although there was a noticeable skew toward LTT compared to the more balanced LTT/sT expression in WaGa cells (Figure 3B, 3C). We were unable to detect expression of sT protein in the MDF cell lines or MCC cell lines, although this issue has been reported on previously and has been attributed to low endogenous expression(39). Vimentin expression confirmed the mesenchymal identity of the transformed cells (Figure 1C), highlighting a distinction from human MCC. However, RNA-seq analysis of the clonal MDF-TA cell lines, MDF-TA-4.1 and MDF-TA-12, indicated a negative enrichment score in EMT-related genes, suggesting a partial shift towards an epithelial transcriptional program following MCPyV TA expression (Figure 3A). Dermal fibroblasts comprise several distinct subtypes with different biological properties, and the susceptibility of dermal fibroblasts to MCPyV-mediated transformation may therefore not be uniform across all dermal fibroblast subtypes(40, 41). Consistent with this possibility, several neuroendocrine markers were detected at low levels in the vector control cells, suggesting that some fibroblast heterogeneity is retained *in vitro* (Figure 2C, 2E, 2F). Accordingly, we cannot determine whether TA-expression directly induces neuroendocrine gene expression or whether fibroblast subpopulations with pre-existing neuroendocrine features are preferentially transformed by MCPyV.

Transcriptomic analysis of the TA-expressing MDFs revealed activation of proliferation-related pathways associated with MCC. Transcription of E2F targets was the most enriched gene set in all TA-expressing MDFs when compared to the vector control cells (Figure 3A), consistent with LTT-mediated inhibition of Rb and activation of E2F-dependent transcription(16). Enrichment of the G2/M checkpoint gene set was also observed in all MDF-TA cells, reflecting their highly proliferative phenotype (Figure 3A). MYC target enrichment was observed in the two clonal MDF-TA cell lines, consistent with the known ability of sT to interact with MYCL and promote oncogenesis by altering transcriptional regulation in MCC(18). In contrast, although aberrant mTOR signaling is a feature of MCC, increased mTOR pathway activity was observed only in MDF-TA-12 cells compared with the vector control(12, 36, 42). Interestingly, Rb appeared to be inhibited in all TA-expressing cells, whereas p53 remained functional (Figure 3D, 3F). This likely reflects the LTT-dominant balance of T antigen expression in our model, as LTT has previously been shown to activate p53, while sT suppresses p53 function(19).

We successfully established tumors using the MDF-TA-12 cells in both NSG and C57BL/6 mice (Figure 4A). Histological analysis revealed highly cellular, poorly differentiated tumors (Figure 4C). In NSG mice, tumor cells infiltrated nearby muscle tissue, indicating aggressive growth (Figure S5B). Tumor tissue exhibited uniform LTT expression (Figure 4G), but expression of CK20 was more heterogeneous, with some cells exhibiting the characteristic perinuclear dot-like pattern (Figure 4G). Although tumors grew progressively in NSG mice, they regressed in C57BL/6 mice after approximately one week, indicating that the transformed cells alone were insufficient to overcome immune surveillance in an immunocompetent host.

MCPyV+ MCC is considered a highly immunogenic tumor owing to expression of the MCPyV TAs, yet successful tumors acquire multiple mechanisms to evade immune surveillance (43). Downregulation of MHC-1 expression is frequently observed in MCC, allowing tumor cells to escape recognition by cytotoxic T cells(44). STING is also silenced in MCPyV+ MCC and may contribute to immune evasion(45). In some cases, MCC tumors express the checkpoint protein PD-L1, which contributes to T-cell dysfunction and exhaustion(5, 6). Importantly, these immune-evasive mechanisms have not been shown to arise as a direct result of MCPyV TA expression and may instead reflect adaptations acquired during tumor evolution under immune pressure. Consistent with this idea, transformation of fibroblasts *in vitro* may lack the necessary selective pressure to induce the immune-evasive changes associated with MCPyV+ MCC. To determine whether the MDF-TA-12 cells could acquire these adaptations, we subjected the cells to serial *in vivo* passaging in immunocompetent C57BL/6 mice. While we were able to achieve improved tumor growth and delayed tumor clearance, the tumors ultimately regressed. Serial *in vivo* selection resulted in decreased MHC-1 expression and reduced expression of p53 target genes (Figure 5C-E), indicating that immune pressure promotes adaptive remodeling of multiple immune-related pathways. Although we hypothesized that *in vivo* selection pressure might upregulate PD-L1 in the tumor cells, unexpectedly we observed significant decreases in PD-L1 expression following serial *in vivo* passage (Figure 5F). This finding suggests that immune pressure does not simply select for a single immune-evasion mechanism but instead drives broader remodeling of the tumor state.

One of the most interesting findings of this study was the coordinated shift from LTT towards sT expression during serial *in vivo* passaging (Figure 5A, 5B). Rather than simply selecting for reduced antigen presentation, our findings suggest that immune pressure may also reshape viral oncogene expression. This observation is particularly intriguing because sT is considered the dominant transforming oncogene in MCPyV-positive MCC and is required for maintenance of the transformed phenotype(12). Moreover, LTT and sT have distinct functions in MCC biology, including opposing effects on p53 signaling and inflammatory responses(19, 20), while sT is also known to stabilize LTT(46). Immune pressure during serial *in vivo* passaging may favor sT expression, and it seems plausible that immunoediting occurring during MCC oncogenesis may influence splicing of the T antigen transcript to fine tune the relative abundance of each isoform and achieve an optimal balance of the two oncogenes. Notably, decreases in MHC-1 expression and p53 target gene expression preceded detectable changes in LTT/sT expression (Figure 5), indicating that immune selection drives coordinated remodeling of multiple pathways, including antigen presentation, p53 signaling, and viral oncogene expression, rather than selecting for a single immune-evasion mechanism. Importantly, some adaptations seen in this model are consistent with adaptations thought to occur during MCC progression and highlight the value of this immunocompetent model for studying tumor evolution under immune pressure. In summary, our findings establish a unique experimental platform for dissecting how MCPyV-driven tumors evolve in the face of immune pressure and identifying mechanisms of immune evasion that may ultimately inform therapeutic strategies.

## Acknowledgments

We thank the members of our laboratories for their insightful discussions and continued support. This work was supported by the National Institutes of Health through grants R01CA187718 and R01CA284690 awarded to J.Y., as well as by the National Cancer Institute through grant P01CA281867 (Project 3) to J.Y. J.R. is supported by NCI grant T32CA288356. The funders had no role in study design, data collection, and interpretation or the decision to submit the work for publication.

---

**Figure S1.**
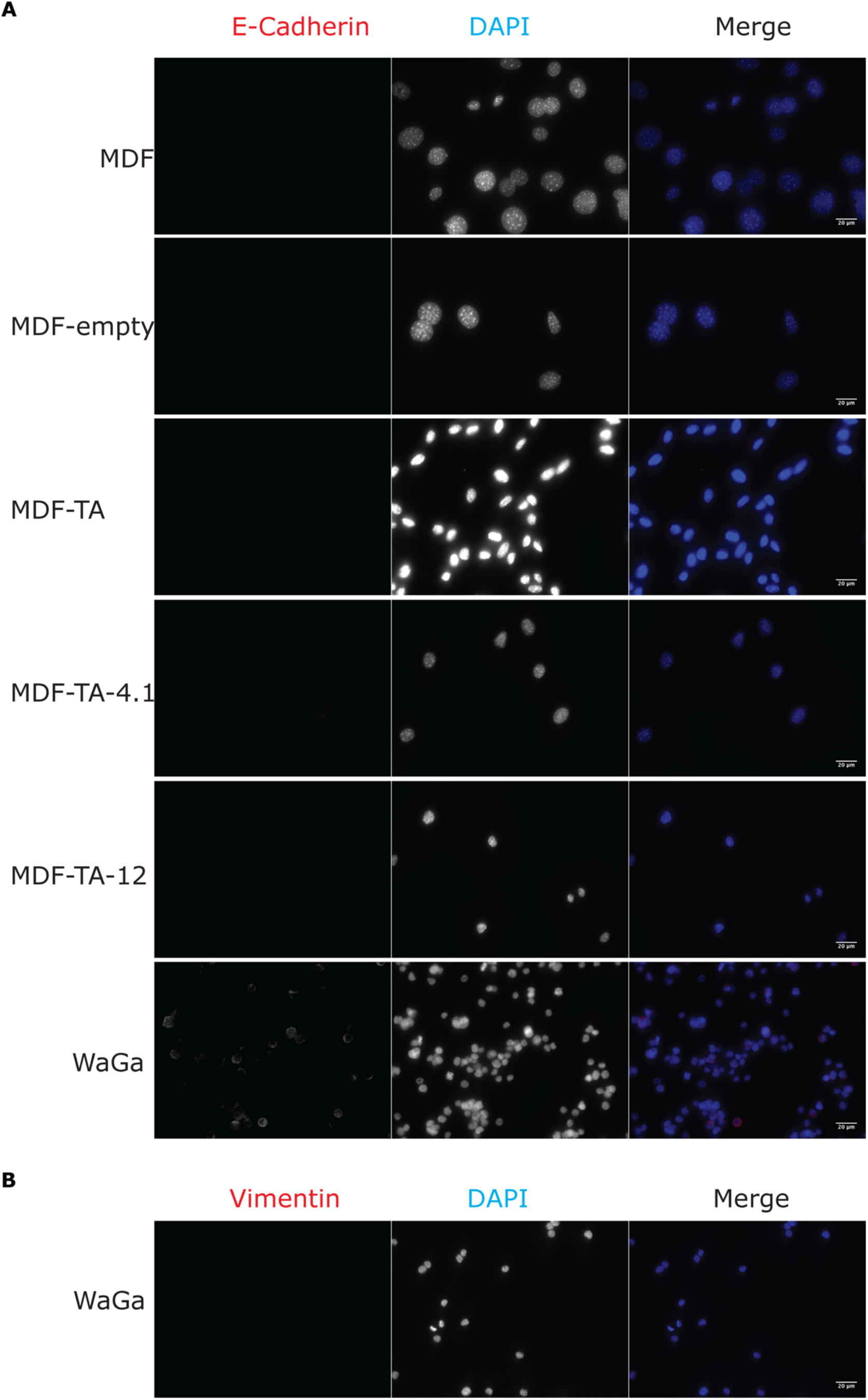
Validation of MDF identity. (A) E-cadherin staining of MDFs, MDF-empty, MDF-TA, MDF-TA-4.1, MDF-TA-12, and WaGa cells at 40X magnification. (B) Vimentin staining of WaGa cells at 40X magnification.

**Figure S2.**
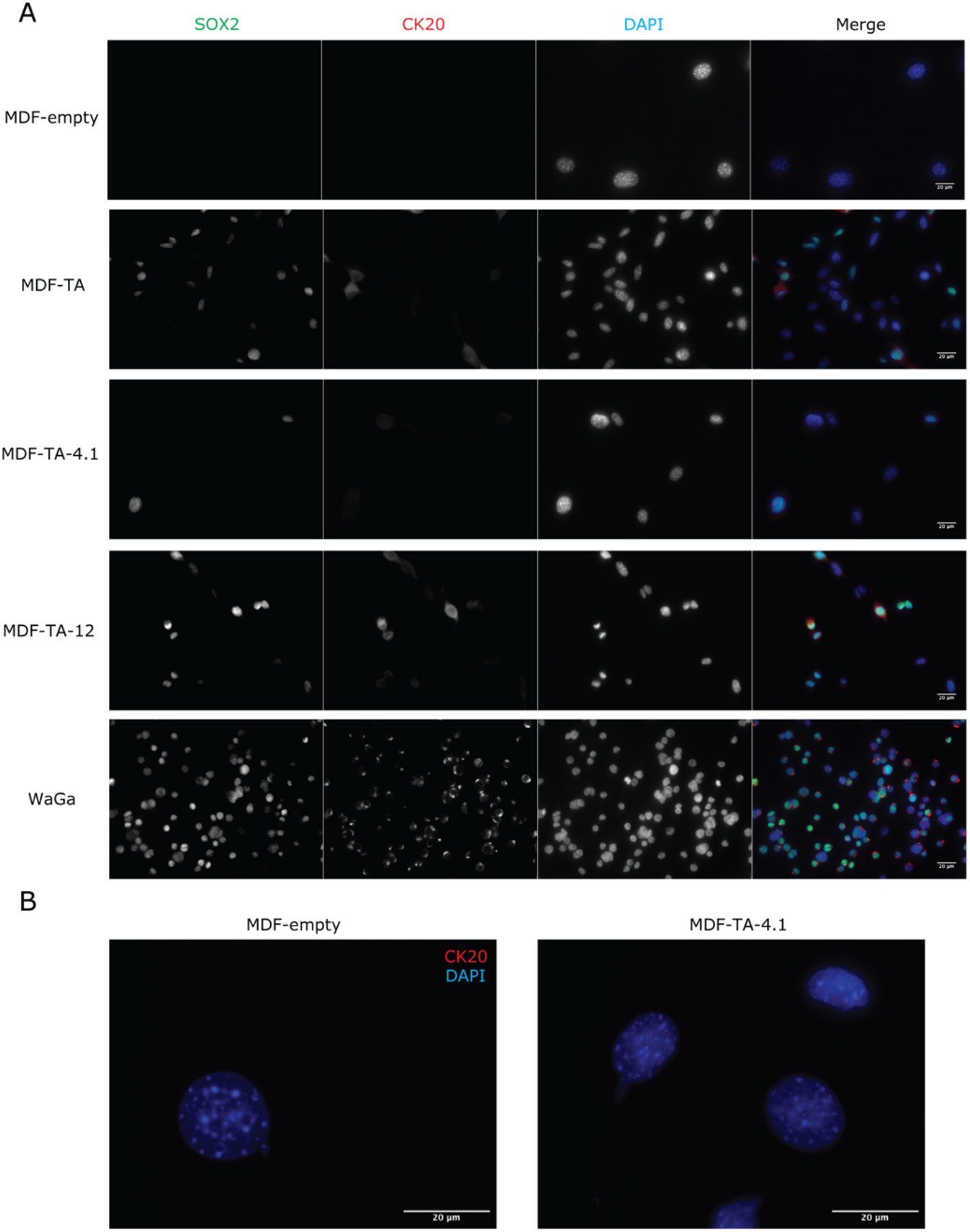
MCC marker expression in MDF cell lines. (A) Immunofluorescent staining of MCC markers SOX2 and CK20 in MDF-empty, MDF-TA, MDF-TA-4.1, MDF-TA-12, and WaGa cells. Images were acquired at 40X magnification under identical conditions. (B) Immunofluorescent staining of CK20 in MDF-empty and MDF-TA-4.1 cells imaged at 100X magnification.

**Figure S3.**
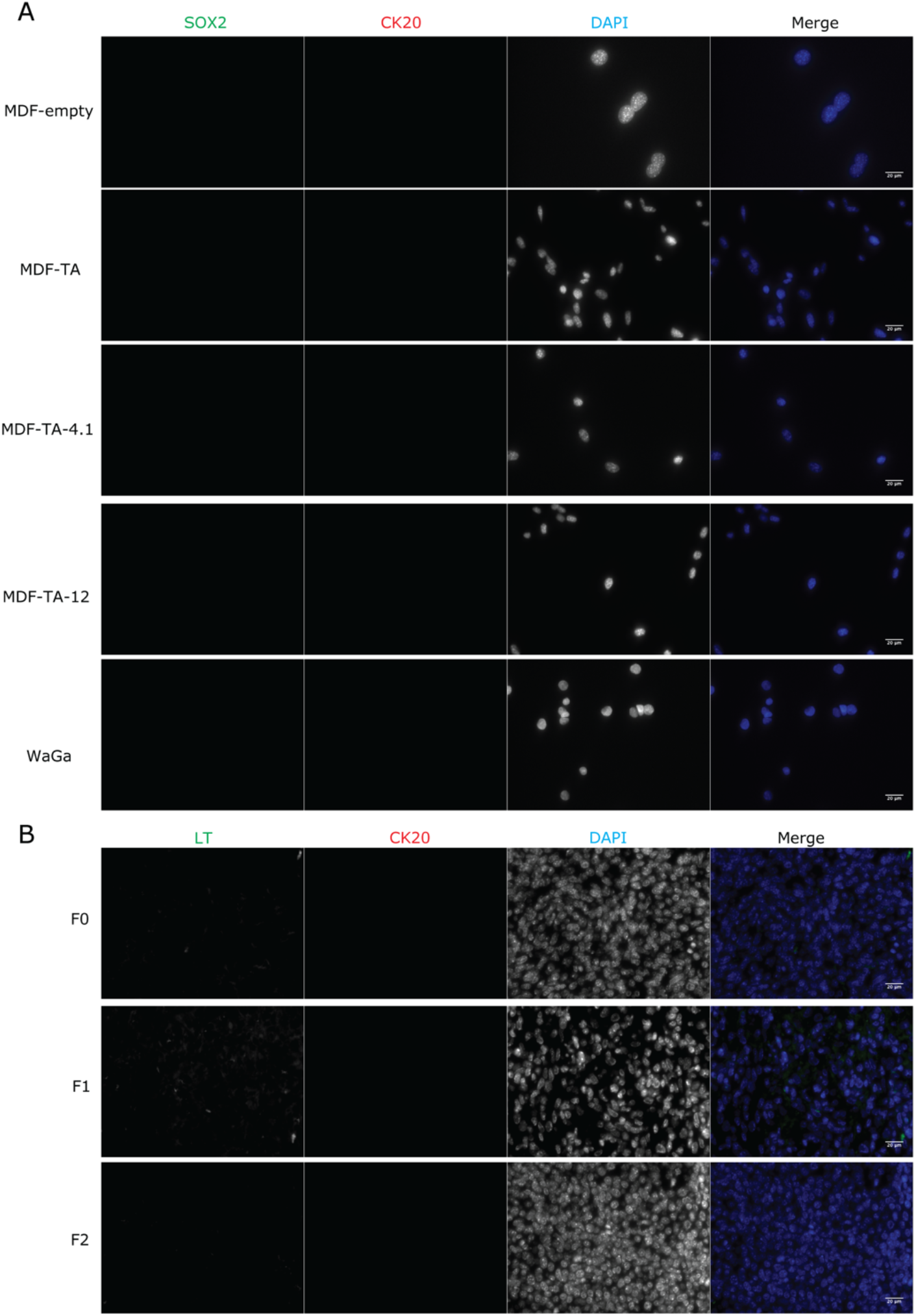
Secondary-only controls for immunofluorescent staining. (A) Secondary only controls for SOX2 and CK20 staining shown in Figure 2 and Figure S2. (B) Secondary only controls for LT and CK20 staining shown in Figure 4.

**Figure S4.**
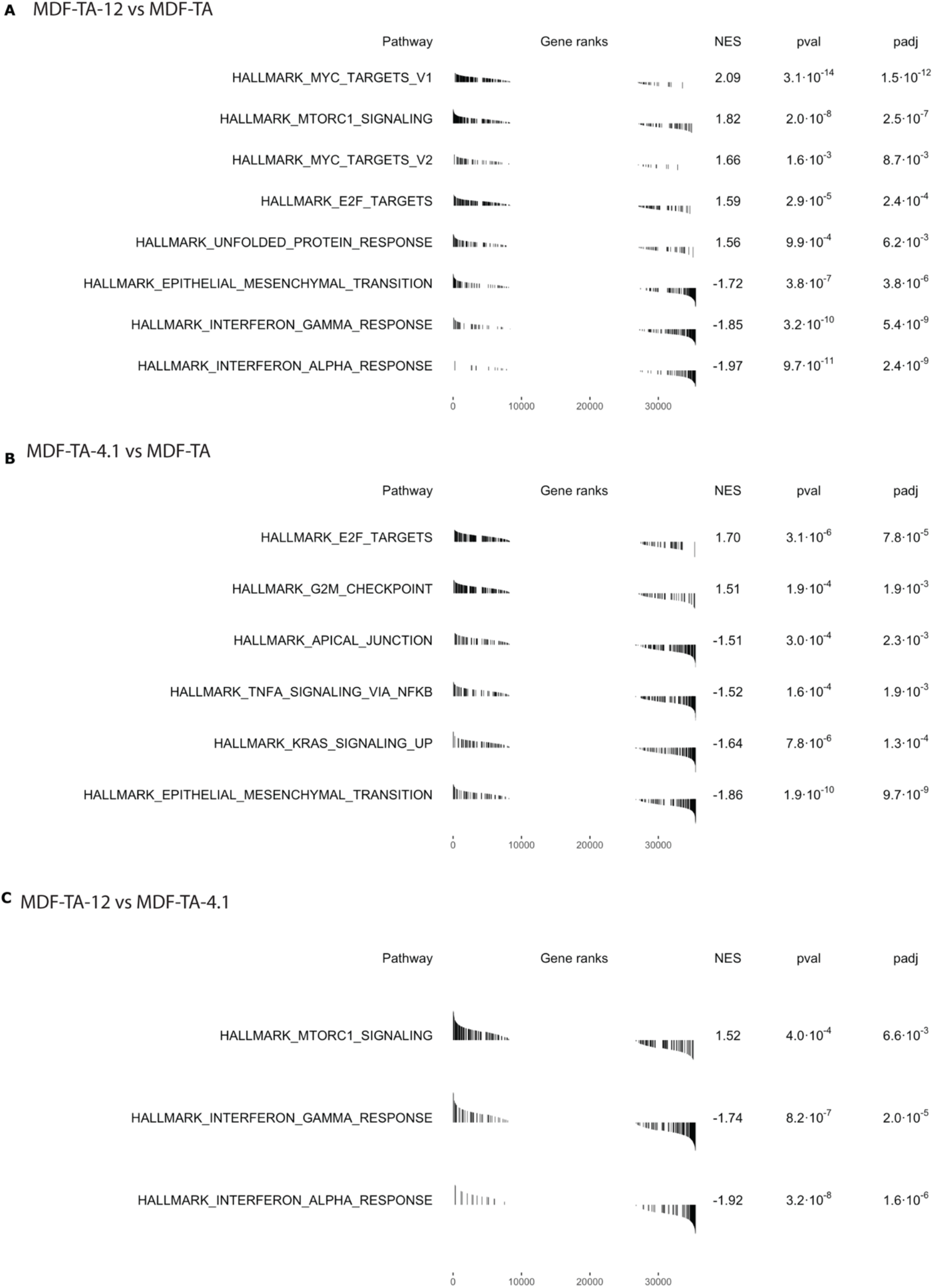
FGSEA analysis of mSigDB hallmark gene expression in MDF-TA cell lines. (A) Comparison of MDF-TA-12 to MDF-TA (B) Comparison of MDF-TA-4.1 to MDF-TA (C) Comparison of MDF-TA-12 to MDF-TA-4.1. All gene sets with a net enrichment score (NES) >1.5 or <-1.5 and an adjust p-value (padj) < 0.01 are shown.

**Figure S5.**
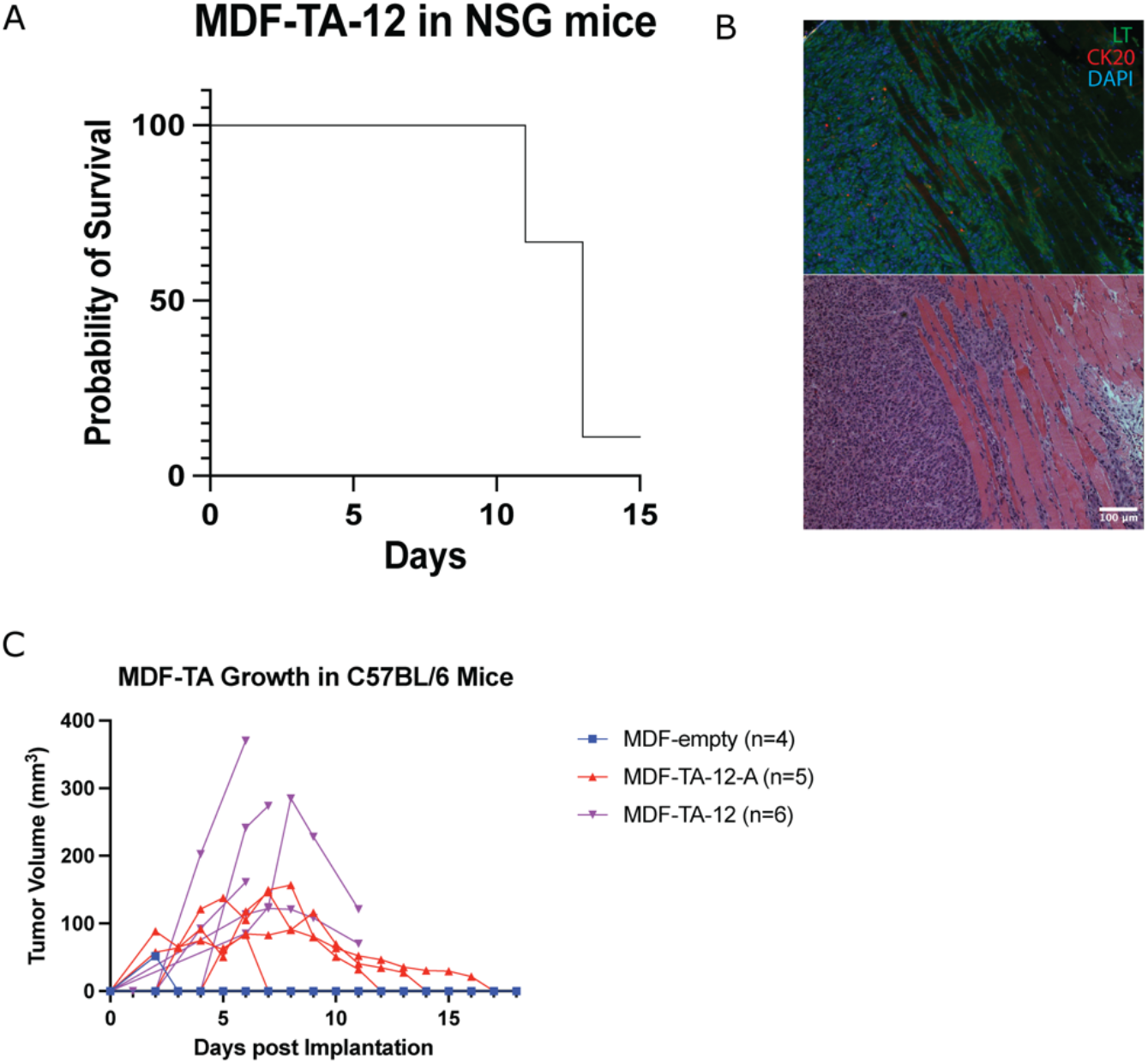
MDF-TA-12 grows aggressively in NSG mice. (A) Survival curve of NSG mice that received MDF-TA-12 cells subcutaneously (n=9). Three mice were euthanized 11 days following tumor cell injection with two mice having impaired mobility and one mouse having a body condition score of 1. Five mice were euthanized 13 days following tumor cell injection due to severe lethargy, ruffled fur, and hunched posture. (B) Invasion of muscle tissue by MDF-TA-12 cells. The same area of the tumor is shown with LT and CK20 staining (top) and hematoxylin and eosin staining (bottom). (C) Growth curves of MDF-TA-12-A cells (isolated from ascites fluid in an NSG mouse) and MDF-TA-12 F0 cells.

**Figure S6.**
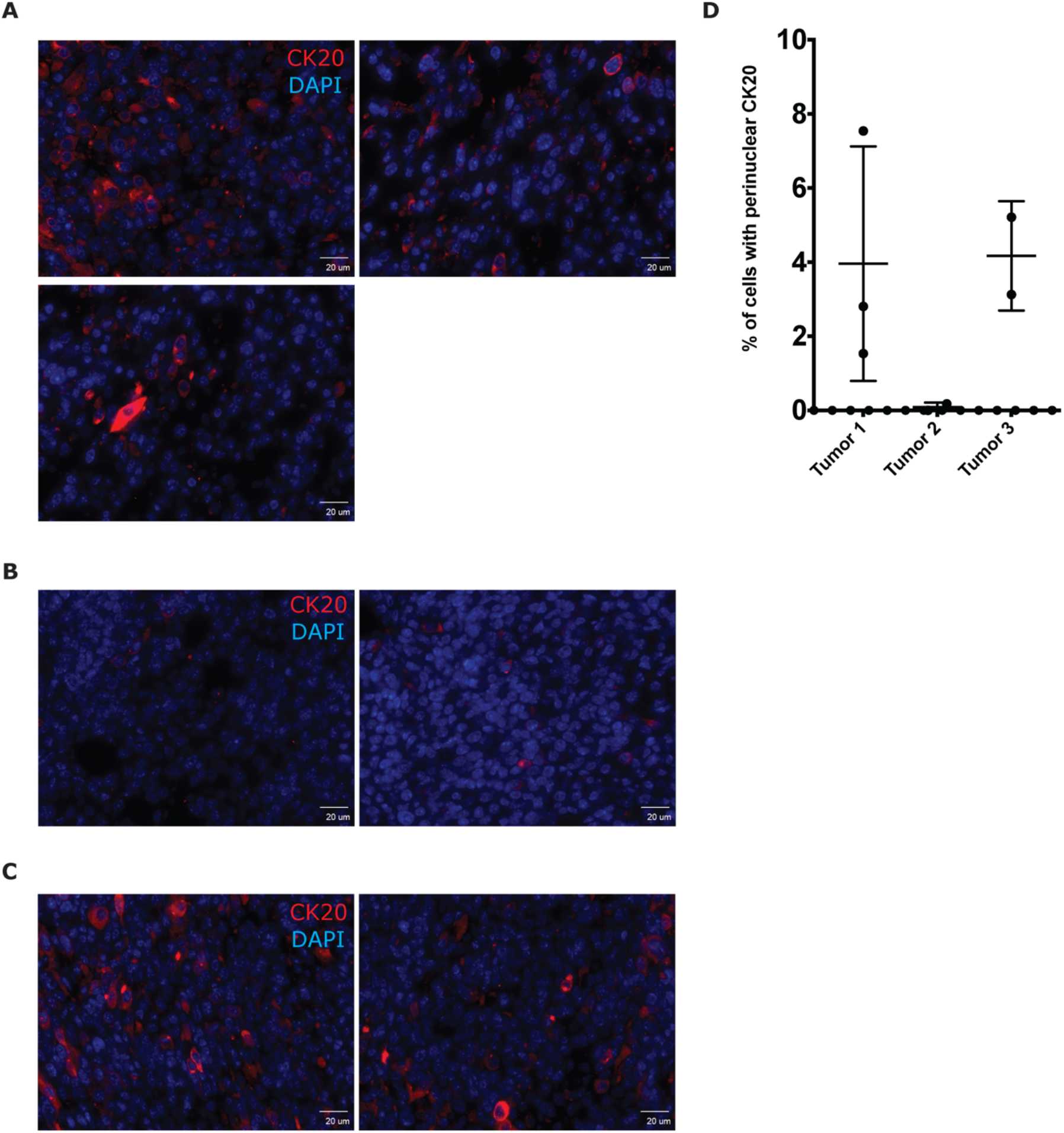
Quantification of CK20 positivity in F0 Tumor tissue. (A) CK20 staining in F0 tumor 1 imaged at 40X magnification. (B) CK20 staining in F0 tumor 2 imaged at 40X magnification. (C) CK20 staining in F0 tumor 3 imaged at 40X magnification. (D) Quantification of cells with CK20 expression in F0 tumors

